# RNA foundation models enable generalizable endometriosis disease classification and stable gene-level interpretation

**DOI:** 10.64898/2026.02.24.707424

**Authors:** Niccolo McConnell, Jennifer Kelly, Ravikanth Tadikonda, Natalia Mulligan, Joao Bettencourt-Silva, Matthew Madgwick, Ritesh Krishna, James Strudwick, Ashley Evans, Stephen Checkley, Anna Paola Carrieri, Michalis Smyrnakis, Charles H Knowles, Laura-Jayne Gardiner

## Abstract

Endometriosis is a chronic inflammatory condition with significant diagnostic delays impacting one in ten reproductive age women worldwide. While machine learning (ML) models trained on transcriptomic data show promise for disease prediction, limited generalizability across independent patient cohorts has hindered clinical translation. Foundations models (FMs) pretrained on large-scale transcriptomic data offer promise to learn transferrable, biologically meaningful representations that could support cross-cohort predictions. We assembled a 12-cohort bulk RNA-seq benchmark (334 samples) and developed a computationally efficient pipeline to test whether FMs improve endometriosis classification, an approach not previously applied to this disease. Using AutoXAI4Omics with cohort-aware validation, we compared embeddings derived from five state-of-the-art RNA FMs against TPM baselines. In cross-cohort prediction, FM embeddings significantly improved performance, achieving a weighted F1-score of 0.83 vs. 0.68 for the baseline. To allow gene-level interpretation of FM embedding models, we introduce classified-aligned integrated gradients (CA-IG), an interpretability approach aligning gene-level attributions to the downstream classifier without end-to-end finetuning. CA-IG revealed a conserved set of predictive genes from FM embeddings across cohort-validation regimes, contrasting with unstable baseline explainability, suggesting that FM embeddings prioritized transferable disease-related signal over cohort-specific effects. These genes include novel candidates that converge on biologically plausible pathways for endometriosis.

## Background

Endometriosis remains a diagnostic and therapeutic challenge, affecting one in ten women of reproductive age and imposing a significant personal and economic burden. Definitive diagnosis still hinges on laparoscopy and biopsy with expected diagnosis times of approximately 9 years in the UK (1–3). The disease is characterized by ectopic endometrial-like tissue and typically presents with chronic pelvic or abdominal pain, dysmenorrhoea, dyspareunia, fatigue, and gastrointestinal or urinary symptoms (4). Endometriosis is also a leading cause of subfertility/infertility, with a prevalence of 30-50% among women seeking infertility care (5). Health-related quality of life is markedly reduced, with increased anxiety and depression; indirect and direct costs are considerable, and in the UK alone the annual economic burden has been estimated at £8.2 billion (6). Clinical assessment is further complicated by phenotypic heterogeneity and a poor correlation between lesion burden and symptom severity, while imaging is often inconclusive for superficial disease and reliable blood or tissue biomarkers remain lacking (7–9). Hindering diagnostic development is the fact that precise molecular drivers of endometriosis physiology are not well defined or understood. As such, there is an increasing focus and need to interrogate high-throughput omics data like transcriptomics, which offer the ability to study genome-wide expression patterns and reveal dysregulated pathways and genes across different patient groups, lesion types and tissue/fluid types. Insights from these types of studies offer potential for improved diagnostics, biomarker identification, disease subtype classification, increased mechanistic understanding of pathogenesis and personalized medicine approaches. This has motivated the application of AI to predict or model endometriosis from high-throughput expression data, including microarrays and RNA-seq, for various applications such as disease classification and biomarker discovery (9–13).

Akter et al. used an ensemble-based approach for endometriosis prediction on single-centre bulk RNA-seq data (n=38) and DNA-methylation data, and reported near-perfect internal performance (F1 ∼ 0.97) using leave-one-out cross- validation (CV) (12). Revisiting the same cohort, another study used elastic-net feature selection with boosted and bagged trees and achieved performance around F1∼0.86 (5-fold CV) (11). On microarrays, Zhang *et al*. (2023) proposed a five-gene panel (*FOS, EPHX1, DLGAP5, PCSK5, ADAT1*) for endometriosis classification and achieved a AUC of 0.836 on a held-out test set and > 0.78 on two external validations (10). A parallel line of work has pursued circulating microRNAs as non-invasive liquid-biopsy markers of endometriosis, showing a six-miRNA panel exhibited a high accuracy for endometriosis prediction (AUC=0.94) in an independent 48-patient validation cohort (9). More recently, Bendifallah *et al*. profiled the circulating miRNome in a prospective 200-patient study and derived an 86-miRNA signature with excellent internal discrimination (AUC∼0.98 under cross-validation), but without multi-centre external replication (13). As the authors note, the broader blood-biomarker literature has been constrained by small cohorts and methodological heterogeneity, yielding few stable, clinically adopted markers. This finding was enforced by Shi *et al.* who demonstrated that subtype-specific tissue markers for deep infiltrating endometriosis were sensitive to cohort composition (14).

While these studies revealed critical biological insights and significantly advanced endometriosis disease understanding, their reliance on single cohorts has limited the generalizability of their models and associated findings to unseen datasets or cohorts. Moreover, this issue limits the clinical applicability and value of their findings in real-world diagnostic settings. As such, there is a need to develop methods or approaches which can derive more generalized insights for endometriosis that will be applicable across a wider range of cohorts and can underpin development of robust models and derivative disease biomarkers. A potential solution to this limitation may be the use of foundation models (FMs), large neural encoders pretrained on vast unlabeled corpora via self-supervision to learn general-purpose representations that transfer across tasks. Recent work has demonstrated the potential of FMs to learn representations that are highly transferrable across different domains, datasets and tasks. In transcriptomics, RNA FMs such as Geneformer (15), scGPT (16), scFoundation (17), and BulkRNABERT (18) are trained on millions of bulk or single-cell gene expression profiles and have shown utility for downstream problems including cell-type annotation, perturbation prediction, and disease classification. Relative to models trained from scratch, these encoders typically yield increased downstream task performance and greater robustness under distribution shift by providing richer, structure-aware features. This study represents the first systematic evaluation of RNA foundation models for endometriosis prediction.

In this work we aimed to investigate if use of RNA FMs meaningfully improved generalizability of the resulting models, allowing accurate cross-cohort disease prediction. To do this we developed a sophisticated cross-dataset evaluation pipeline to test the domain generalizability of FM embeddings using five state-of-the-art RNA FMs; Geneformer (15), scFoundation (17), scGPT (16), BulkRNABERT (18) and BMFM-RNA (19) and a frozen-encoder paradigm (pretrained RNA encoders are kept fixed and used to compute embeddings which were then used as input features for a classical ML task). The resultant pipeline is detailed in Figure 1 and consisted of 1) identification of Gene Expression Omnibus (GEO) ids for relevant bulk RNA-seq endometriosis/control datasets and automated download, extraction, amalgamation and normalization of transcriptomic datasets given these pre-defined ids, and LLM-powered metadata curation with human validation to ensure consistent and high-quality target label generation; 2) extraction of embeddings from pre-trained frozen RNA FMs via one forward pass through each model; 3) generation of robust cohort-aware multi-fold strategy to evaluate within vs. cross cohort learning capabilities of using FM embeddings as features compared to baseline normalized (TPM) gene expression counts; 4) an explainable ML workflow to train and test a decision tree classifier to predict endometriosis status (see Methods) and to evaluate performance of our feature sets; 5) classifier-aligned integrated gradients (IG) method to calculate gene attribution scores and 6) an LLM-powered knowledge graph (KG) approach to interpret biological significance of the most predictive features.

**Figure 1.**
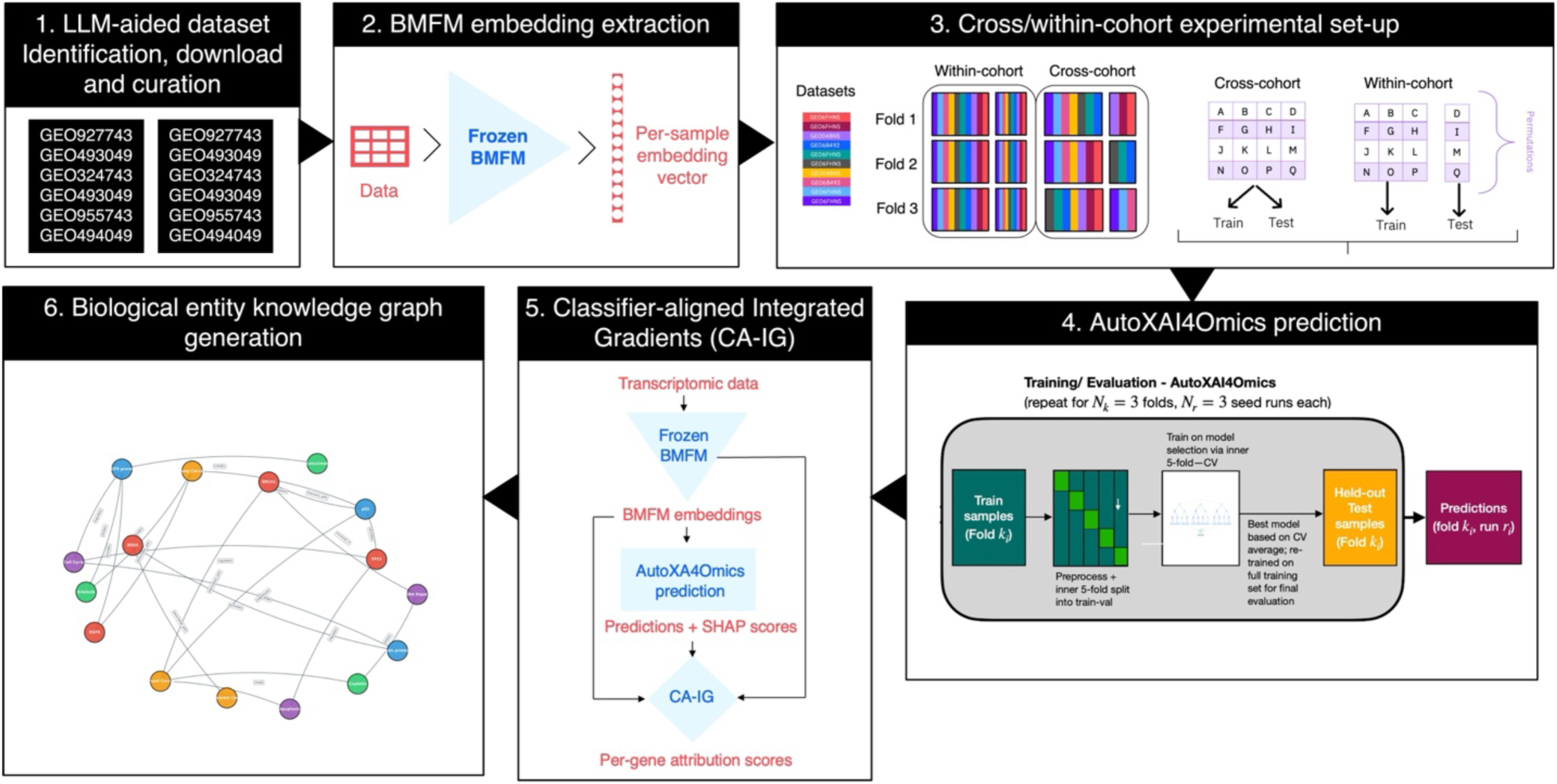
End to end pipeline. Pipeline consists of 1) identification of GEO ids for relevant endometriosis/control datasets and LLM-powered metadata curation with human validation to ensure consistent and high-quality target label generation; 2) extraction of embeddings from pre-trained frozen RNA FMs; 3) generation of robust cohort-aware multi-fold strategy to test within vs. cross cohort learning capabilities of feature sets; 4) an explainable ML workflow to evaluate endometriosis classification performance of our feature sets; 5) classifier-aligned integrated gradients (CA-IG) method to calculate gene attribution scores and 6) an LLM-powered knowledge graph (KG) approach to interpret biological significance of the most predictive features.

Our dataset incorporated a multi-cohort bulk RNA-seq benchmark spanning 12 independent GEO studies (334 samples), one of the largest collections of independent cohorts for bulk RNA-seq endometriosis classification to date, and the first multi-cohort evaluation of frozen RNA foundation-model embeddings for this task. This approach allowed our data to be enhanced with the prior biological knowledge of the FM, without the computational expense of pre-training or fine-tuning. For interpretation of the embeddings derived from the FMs, we applied a hybrid classifier-aligned integrated gradients (CA-IG) method, which can produce stable, biologically plausible gene-level attributions across evaluation regimes. To identify unexplored relationships between our most explainable genes and endometriosis in a sparsely characterized research area, we applied a literature-driven knowledge graph and LLM-based inference method to systematically integrate and reason over published evidence, yielding a literature-grounded representation of gene-endometriosis relationships suitable for hypothesis generation.

This work enabled a robust assessment of the capability of FM-based representations to generalize across heterogenous sources, a persistent challenge situation in computational biology, with often limited availability of large, harmonized datasets. Many studies rely on small sample sizes collected under specific conditions, which commonly leads to individual models with good performance but poor generalizability. This lack of generalizability is of particular concern with understudied complex diseases like endometriosis. By using FM-derived embeddings in this way we aim to address this fundamental issue and understand if representations learnt from pre-trained foundation models capture biologically genuine patterns which transcend dataset-specific biases.

## Results & Discussion

### Novel pipeline design enables evaluation of cross-cohort generalizability

In this work we developed a comprehensive end-to-end pipeline enabling inference-only use of pre-trained RNA foundation models for reproducible endometriosis disease prediction and gene-level interpretation from bulk RNA-seq data (Fig 1). This pipeline integrates: (i) automated download and harmonization of GEO RNA-seq studies with LLM-assisted metadata curation and expert validation, (ii) extraction of FM embeddings via a single forward pass through the model, (iii) cohort-aware evaluation using within-cohort and cross-cohort strategies, (iv) consistent classical ML training and evaluation using AutoXAI4Omics tookit, (v) explainability using SHAP for TPM models and classifier-aligned integrated gradients (CA-IG) for embedding models and (vi) literature-grounded knowledge-graph generation and interpretation, and pathway analysis. We assembled a multi-cohort benchmark spanning 12 independent Gene Expression Omnibus GEO studies (334 samples; 17,518 unified genes) (20), by searching for human studies of endometrial tissue related to endometriosis and restricting results to expression profiling by high-throughput sequencing, comprising 259 endometriosis cases and 75 controls (Table 1). This multi-study setting captures clinically realistic heterogeneity across laboratories and protocols and enables an accurate assessment of generalization under cohort shift.

**Table 1:** Summary of the cohorts used in this study. Detailing GEO ID’s, sample counts and positive proportions of patients or samples within each cohort with a positive diagnosis of endometriosis.

| Dataset ID | Dataset name (GEO name) | Sample count(n) | Proportion of disease cases |
| --- | --- | --- | --- |
| 1 | GSE134056 | 38 | 0.42 |
| 2 | GSE202571 | 17 | 1 |
| 3 | GSE205494 | 12 | 0.5 |
| 4 | GSE135485 | 58 | 0.93 |
| 5 | GSE212787 | 17 | 0.65 |
| 6 | GSE135640 | 36 | 0.5 |
| 7 | GSE226870--GSE226872 | 53 | 1 |
| 8 | GSE157153 | 66 | 1 |
| 9 | GSE230956 | 8 | 0.5 |
| 10 | GSE171653 | 11 | 1 |
| 11 | GSE190580 | 7 | 0.43 |
| 12 | GSE243157--GSE243158 | 11 | 0 |

From these datasets we derived two feature representations for each sample; (i) gene-level log-transformed TPM values, used as a baseline, and (ii) embeddings extracted from state-of-the-art ‘frozen’ RNA FMs. To assess if FM-derived features provide an advantage over baseline features, we evaluated two classification approaches. In approach 1, we used samples from all 12 cohorts to build the training and test sets, simulating a ‘**within-cohort’** approach (see Methods). Here we expect elevated performance since we do not ask the model to generalize to a completely unseen cohort, but in doing so we risk models learning patterns distinct to each dataset. For approach 2, we designed test and train sets which consisted of distinct cohorts, simulating a ‘**cross-cohort’** approach (see Methods). In this approach we trained the model on samples from specified cohorts and tested exclusively on samples from independent cohorts which were not seen during training. Here we expect a drop-off in performance as the model must generalize across many cohorts. The latter scenario reflects a common challenge in clinical and biological research, where data are typically distributed across many small, independent cohort. By comparing performance across these two strategies, we aimed to determine whether FM-derived embeddings improve robustness and generalization in the presence of cohort heterogeneity, relative to classical gene-level features as a baseline.

### Within-cohort disease prediction

We first evaluated the within-cohort performance (where training and test sets include samples from the same cohort, though not the same patient) for endometriosis disease classification (Table 2). To obtain a robust analysis of model performance variability and to allow statistical comparison, we evaluated performance by performing 3-cross validation folds with 5 random-seed runs per fold per model (15 runs in total per model), and performed hierarchical bootstrapping (10,000 iterations) with re-sampling, allowing calculation (for each metric) of mean, mean effect size, 95% confidence interval (CI) and two-tailed p-values (see Methods).

**Table 2.** Within-cohort results. Predictive performance metrics for endometriosis disease prediction using our within-cohort strategy. Four model performance metrics are provided (weighted-F1, class0-F1, class1-F1 and accuracy) from classification models built using the baseline TPM features and FM-embedding features from five RNA FMs. For GeneFormer the V2-104M-HC model was used and for BMFM-RNA the 110m-mlm-rda-v2 model was used. For each metric, table reports the estimated mean performance over 10,000 bootstrap replicates and corresponding low (2.5^th^) and high (97.5^th^) values from calculating the 95% confidence interval (CI) obtained via nested bootstrapping which re-sampled folds and seed runs with replacement. All metrics are computed on the held-out test set, and ML models were built and evaluated using AutoXAI4Omics ML pipeline (see Methods), using AdaBoost.

| Model Name | Weighted F1 | Class 0 – F1 | Class 1 – F1 | Accuracy |
| --- | --- | --- | --- | --- |

|  | (mean [95%CI]) | (mean [95%CI]) | (mean [95%CI]) | (mean [95%CI]) |
| --- | --- | --- | --- | --- |
| **TPM baseline** | 0.86 [0.84,0.87] | 0.69[0.67, 0.71] | 0.91 [0.90,0.92] | 0.86 [ 0.84, 0.87] |
| **BulkRNABert** | 0.89 [0.88,0.91] | 0.77 [0.75, 0.79] | 0.93 [0.92,0.94] | 0.89 [ 0.88, 0.91] |
| **scFoundation** | 0.85 [0.81, 0.87] | 0.63 [0.55,0.69] | 0.91 [0.89,0.92] | 0.85 [0.82, 0.88] |
| **scGPT** | 0.85 [0.81, 0.87] | 0.65 [0.57,0.72] | 0.91 [0.89,0.93] | 0.86 [0.82, 0.89] |
| **GeneFormer** | 0.90 [0.86, 0.91] | 0.76 [0.69,0.81] | 0.93 [0.91, 0.94] | 0.89 [0.86, 0.91] |
| **BMFM-RNA** | 0.85 [0.81,0.90] | 0.66[0.55, 0.77] | 0.90 [ 0.88, 0.94] | 0.85 [0.70,0.81] |

ML classification of endometriosis disease using the log transformed TPM data (baseline) resulted in a mean weighted-F1 score of 0.86, a class 1-F1 of 0.91 (disease), a class 0-F1 of 0.69 (no disease) and an accuracy of 0.86 (Table 2, Fig 2A), indicating good performance for endometriosis classification from multi-cohort transcriptomic data, with more accurate prediction of disease than healthy samples (Table 2).

**Figure 2:**
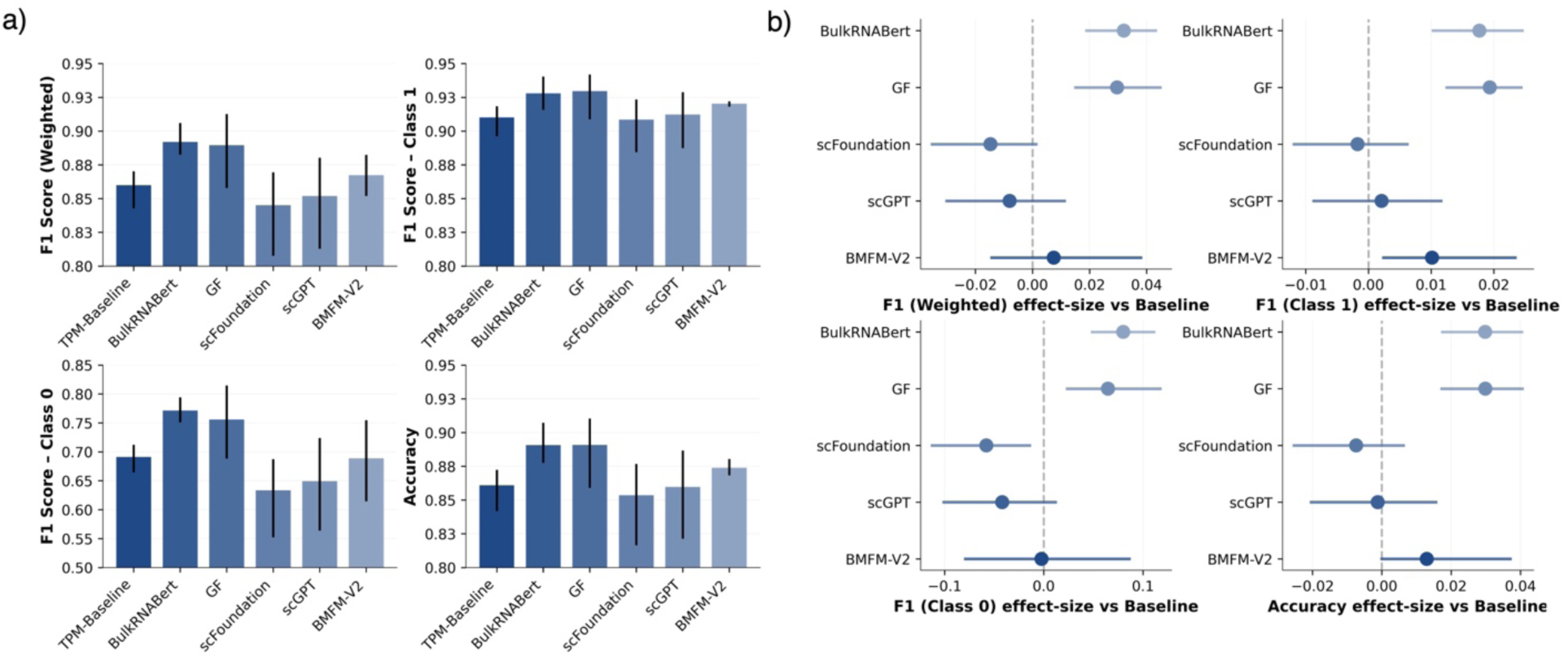
**Within-cohort results**. (a) Mean predictive performance metrics for endometriosis using our within-cohort strategy for the baseline TPM features versus using frozen embeddings as from the five RNA FMs as features. Bars show the mean across outer folds and seeds; whiskers denote 95% CIs. Metrics reported: weighted-F1 (primary), F1 for the positive class (class 1; endometriosis), F1 for the negative class (class 0; controls), and accuracy. (b) Effect size relative to the baseline: mean difference in score (embedding model minus baseline) with 95% Cis. All models use identical training and evaluation within the series-aware within-cohort splits

Next, we computed the statistical significance of pairwise model performance by calculating two-tailed p-values from the bootstrapped samples. Among the five frozen FMs, embeddings extracted from BulkRNABERT and Geneformer surpassed the predictive performance of the TPM baseline classifier in all four metrics with statistical significance (P < 0.05, Supplementary Table S1A), with the weighted F1 increasing from 0.86 for the TPM classifier to 0.89 and 0.90 for BulkRNABERT and Geneformer classifiers respectively (Table 2). Prediction of the healthy class (class 0) was also significantly improved using BulkRNABERT and Geneformer embedding classifiers, increasing from a baseline-F1 of 0.69 to F1 scores of 0.77 and 0.76 for BulkRNABERT and Geneformer classifiers respectively (Table 2, Supplementary Table S2A, Fig 2). For the remaining three FM embedding classifiers (scFoundation, scGPT and BMFM-RNA) we saw reduced or equal model performance compared to the baseline TPM classifier for all metrics, and for prediction of the healthy class (class 0), the scFoundation embedding classifier performed significantly worse than the TPM model (P < 0.05, Supplementary Tables S2A, S3A and S4A). Surprisingly, our top performing FM-embedding classifier was using embeddings extracted from the single-cell pretrained FM Geneformer, whereas we were using bulk RNA-seq data. This suggests that single-cell priors transfer effectively to bulk RNA-seq.

For each FM-derived feature set classifier, mean effect-size (+ 95% CI) was also calculated which indicated the average difference in performance metrics from the raw baseline transcriptomic feature set model, which we treated as our baseline (Fig. 2B, Supplementary Table S5). Although we observed an increase in F1 score and positive F1 effect sizes for two FM-derived embedding classifiers, increases were moderate in scale with very small effect sizes, and we observed a drop in model performance for the remaining three FM embedding classifiers. Overall, within-cohort evaluation yielded high performance for both feature representation type (TPM and embeddings), with only modest and model-dependent benefits from embeddings (Fig 2B). This pattern is consistent with a setting where cohort-specific structure is shared between test and train data, and in these conditions a well-tuned classical pipeline using gene-level features already performs strongly, with embeddings adding limited advantage.

### FM-derived embeddings substantially improve cross-cohort disease prediction

We next evaluated cross-cohort generalization, where entire cohorts are held out for testing. For this approach, we constructed balanced grouped cohort splits with *K*=3 outer folds to match the within-cohort approach (see Methods). Individual cohorts were assigned to test folds by a constrained random search (1,000 trials; fixed seed) that (i) enforced a minimum test size (at least 60 samples per fold) and (ii) minimized the across-fold standard deviation of the positive fraction (disease positive samples) and of fold sizes. We report the resultant test/train dataset assignments in Methods. Mean model performance metrics and effect sizes were calculated as previously described (see Methods).

When using our cross-cohort approach for disease prediction, the baseline classifier (log-transformed TPM) achieved a mean weighted F1-score of 0.68, a marked decline from the F1-score of 0.86 obtained from the within-cohort approach (Table 3). This loss in performance was expected, as the trained model was less able to generalize to samples from unseen cohorts, likely due to differences in sample collection, patient clinical diversity, feature distribution and class-imbalances. All embedding-based classifiers also exhibited a drop in predictive performance when trained and tested using the cross-cohort vs. within-cohort approach, however the degree of reduction was substantially lower in 4/5 of the embedding classifiers compared to the baseline classifier, with drops in mean weighted F1-score of 0.07, 0.07, 0.10 and 0.11 for GeneFormer, scGPT, BMFM-RNA and BulkRNABert models respectively, compared to a drop of 0.18 for the baseline (Table 3).

**Table 3.** Predictive performance metrics from endometriosis disease prediction (cross-cohort). Four model performance metrics are provided (weighted-F1, class0-F1, class1-F1 and accuracy) from models using the baseline TPM features and FM-embedding features from 5 SOTA transcriptomic FMs. For GeneFormer the V2-104M-HC model was used and for BMFM-RNA the 110m-mlm-rda-v2 model was used. For each metric, table reports the estimated mean performance over 10,000 bootstrap replicates and corresponding low (2.5^th^) and high (97.5^th^) values from calculating the 95% confidence interval (CI) obtained via nested bootstrapping which re-sampled folds and seed runs with replacement. All metrics are computed on the held-out test set, and ML models were built and evaluated using AutoXAI4Omics ML pipeline (see Methods), using AdaBoost.

| Model Name | Weighted F1<br>(mean[95%CI]) | Class 0 – F1<br>(mean [95%CI]) | Class 1 – F1<br>(mean [95%CI]) | Accuracy<br>(mean [95%CI]) |
| --- | --- | --- | --- | --- |
| **TPM baseline** | 0.68 [0.59,0.78] | 0.42 [0.23, 0.53] | 0.75 [0.69,0.85] | 0.65 [ 0.57, 0.77] |
| **BulkRNABert** | 0.78 [0.68, 0.90] | 0.53 [0.21, 0.73] | 0.86 [0.82,0.91] | 0.79 [ 0.71, 0.86] |
| **scFoundation** | 0.58 [0.52, 0.63] | 0.31 [0.08,0.41] | 0.66 [0.55,0.79] | 0.57 [0.50,0.66] |
| **scGPT** | 0.78 [0.74, 0.81] | 0.50 [0.32,0.61] | 0.86 [0.85,0.87] | 0.79 [0.77, 0.80] |
| **GeneFormer** | 0.83 [0.77, 0.87] | 0.62 [0.44,0.74] | 0.89 [0.86, 0.91] | 0.83 [0.78,0.87] |
| **BMFM-RNA** | 0.80 [0.71,0.86] | 0.56 [0.29,0.70] | 0.87 [0.83,0.91] | 0.81 [0.73,0.86] |

In contrast to our within-cohort approach, now when using a cross-cohort approach, all FM-embedding classifiers significantly outperformed the TPM-baseline model for disease classification, except for scFoundation (*P <* 0.05, Supplementary Table S1B). Geneformer and BMFM-RNA were the top two performing embedding models for all metrics (mean weighted F1-score, class 0 F1-score, class 1 F1-score and mean accuracy) with mean weighted F1-scores of 0.83 and 0.80 respectively (Table 3, Fig 3, *P <* 0.05). Significant increases in performance for all embedding models apart from scFoundation compared to baseline were also apparent for class 1 F1-score and accuracy metrics (*P <* 0.05), while GeneFormer, BMFM-RNA and scGPT also achieved significantly higher F1-scores for class 0 compared to baseline (*P <* 0.05, Table 3, Supplementary Table S1B). For the outperforming FMs, we saw an average positive effect size (across all four models) compared to the baseline of 0.11 for weighted F1, class 0 F1-score and class1 F1-score and of 0.13 for accuracy (Supplementary Table S6). We saw the greatest performance improvement from Geneformer and BMFM-RNA embeddings, where class 0 F1 increased (compared to TPM baseline) by an average of 0.2 and 0.14 respectively, and the weighted F1-score increased by an average of 0.15 and 0.12 respectively, reflecting a substantial and significant improvement in model performance (Table 3, Fig 3, Supplementary Table S6).

**Figure 3:**
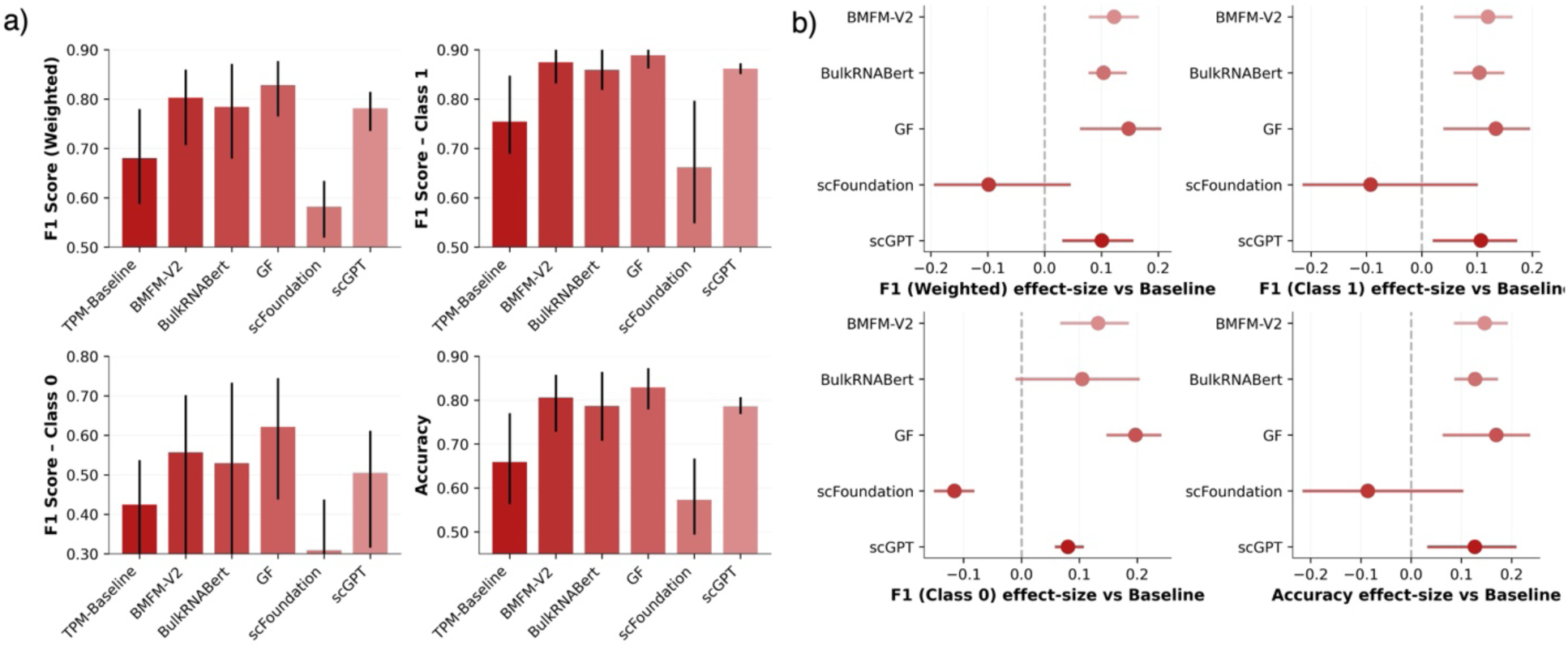
Cross-cohort results (balanced grouped-cohort validation). (a) Mean performance for the (TPM-Baseline) versus the same pipeline using frozen embeddings (BulkRNABERT, scFoundation, scGPT, Geneformer, BMFM-MLM-RDA). Bars show the mean across outer folds and seeds; whiskers denote 95% CIs. Metrics: weighted-F1 (primary), F1 for the positive class (class 1), F1 for the negative class (class 0), and accuracy. (b) Effect size relative to the baseline: mean score difference (embedding model minus baseline) with 95% CIs; the vertical dashed line at 0 marks parity (values > 0 favour embeddings). In this setting, train and test cohorts are disjoint: each outer fold holds out a group of GEO Series (i.e., of cohorts) for testing, and all model selection is confined to the training side

### Gene-level attribution enables interpretation of baseline TPM model

We investigated explainability of the within-cohort and cross-cohort baseline TPM classifiers using SHAP (Shapley Additive Explanations) a model-agnostic attribution framework which computes feature importance scores using game theory (21). This approach allows us to identify the genes most important for disease prediction. Using results from all folds (n=3) and all runs (n=5) per model, for each true positive sample (endometriosis disease positive, correctly predicted as disease) we first extracted SHAP values for all genes and normalized across folds. To focus on disease-driving signal our normalization approach included first setting all negative SHAP scores to zero and normalizing the remaining positive samples, resulting in proportionate importance scores for disease-driving genes. Normalized SHAP values were then averaged across models, runs and folds, resulting in one average SHAP value per gene per model. For the within-cohort baseline TPM model (AdaBoost) across all runs and folds there were an average of 75 true positive samples from which the SHAP values were normalized and averaged. A total of 51 genes exhibited non-zero SHAP values, with the top 20 disease-associated genes shown in Table 4. For the cross-cohort baseline TPM model (AdaBoost) across all runs and folds there were an average of 111 true positive samples from which the SHAP values were normalized and averaged with a total of 64 genes that exhibited non-zero SHAP values (Table 4).

**Table 4.**
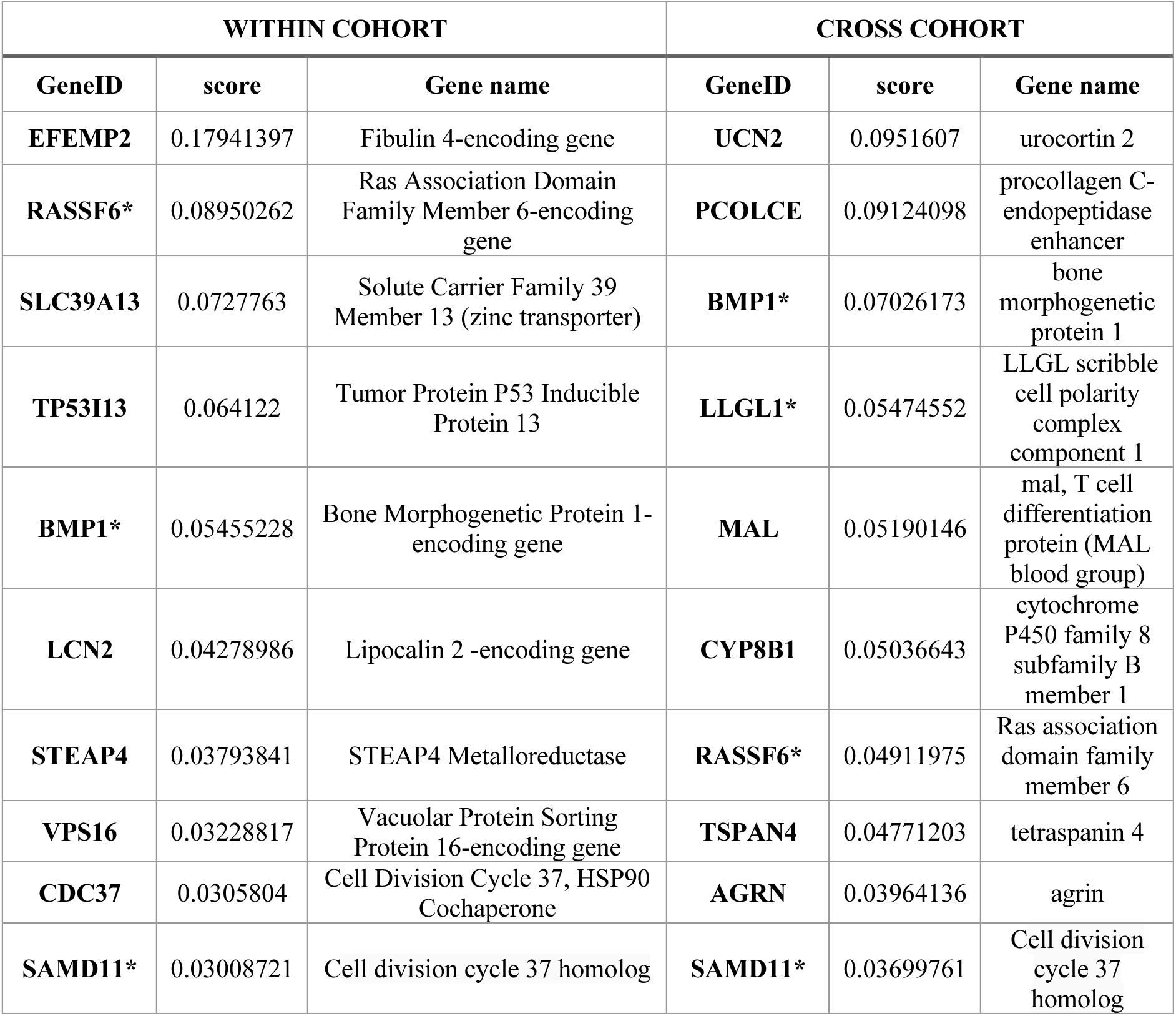
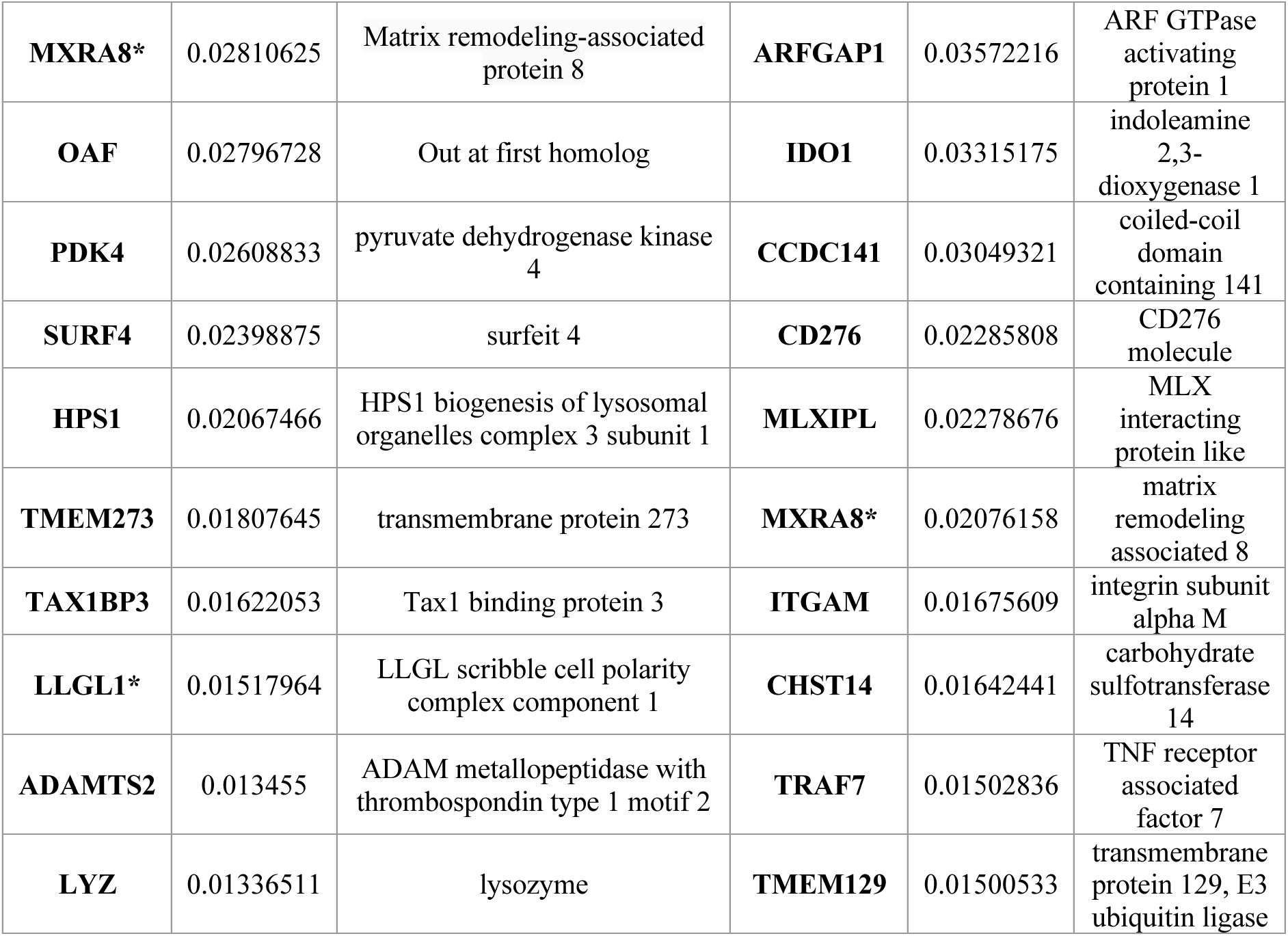
Baseline TPM explainability. Top 20 positive (disease-driving) genes identified via SHAP for TPM baseline model for the within-cohort approach and compared to the cross-cohort approach. Normalized using a positive-only approach, then averaged across runs and folds (HGNC symbols) (Full gene list available in Supplementary Tables S7 (within-cohort) and S8 (cross-cohort)).

We observed little overlap between the TPM-baseline explainability results for the within-cohort vs cross-cohort approaches, with only 5/20 of the top 20 explainable genes in the within-cohort approach also within the top 20 for the cross-cohort approach (Table 4). The top four most important genes for the within-cohort and cross-cohort approaches were completely distinct, being derived from different biological pathways (WikiPathways), indicating that different biological signals were being captured by the within and cross-cohort approaches (Supplementary Figure 1, Table 4). This variability in the predictive features between models is concerning for researchers using interpretation of ML models to guide biomarker or target discovery for disease and could partially explain our drop in performance for cross-cohort analysis if the predictive features from the within-cohort performance do not generalize in the cross-cohort context i.e., cohort specific predictors are being learned in the within-cohort context.

The five overlapping genes between the within and cross-cohort approaches were *RASSF6*, *BMP1*, *MXRA8*, *SAMD11* and *LLGL1*. *RASSF6* encodes a Ras-effector protein which binds activated Ras and is involved in induction of apoptosis. It’s a member of a tumor repressor network which is highly conserved, and it’s inactivation or suppression has been linked to several diseases including cancer (22). *BMP1* (bone morphogenic protein 1) is a metalloprotease which regulates formation of extracellular matrix (ECM) by cleaving/processing of various precursor proteins. Mutations in this gene have been associated with skeletal/connective tissue disorders (23). The role of *MXRA8* (Matrix Remodeling Associated 8- encoding gene) is less established, but it is thought to be involved in glial blood-brain barrier functioning and has been associated with glioma progression and a suggested therapeutic target for glioma (24). *MXRA8* is known to play roles in inflammation and cell signaling that could be potentially relevant to endometriosis. Long non-coding RNA transcribed from the *SAMD11* region has been functionally linked to endometrial biology (decidualization) (25). *LLGL1* is a well-known tumor suppressor gene involved in maintaining epithelial cell polarity and tissue architecture in various cancers, including endometrial cancer (26).

### Classifier-aligned integrated gradients (CA-IG) allows FM embedding gene–level interpretation

Next, we sought to understand if and how model explainability differed when using FM embeddings for disease classification. Because the downstream classifier is not jointly fine-tuned with the encoder, standard gradient attribution cannot be directly applied. We therefore developed classifier-aligned integrated gradients (CA-IG), an attribution approach that assigns importance to FM gene tokens by computing Integrated Gradients (IG) through the frozen encoder along a classifier-relevant direction in the embedding space. This direction is defined locally for each sample using SHAP scores from the downstream classifier with respect to the FM embedding, ensuring that gene-level attributions reflect the features used by the classifier. Importantly, SHAP is used solely to define this classified-aligned weighting of embedding dimensions, while IG performs the attribution itself. This formulation avoids backpropagation through the non-differentiable classifier, substantially reducing computational cost while preserving biological interpretability by providing gene-level explanations (see Methods).

Using our CA-IG approach to extract explainability for the FM-embedding classifiers, we observed a striking conservation of disease-associated genes between the within-cohort and cross-cohort settings (Table 5). Of the top 20 genes identified in the within-cohort analysis, 18 overlapped within the cross-cohort top 20, and conversely 18/20 of the top cross-cohort genes were also present in the within-cohort list (Table 5). Notably, the top five genes were identical and ranked in the same order for both strategies, indicating highly consistent relative importance across evaluation regimes (Fig 4). This contrasts sharply with the TPM-based models, where predictive features varied substantially between within and cross cohort analyses. The stability of the embedding-derived attributions parallels the reduced performance drop observed for the FM-based classifiers under cohort shift, suggesting that the pretrained representations may prioritise transferable disease-related signal over cohort-specific effects.

**Figure 4:**
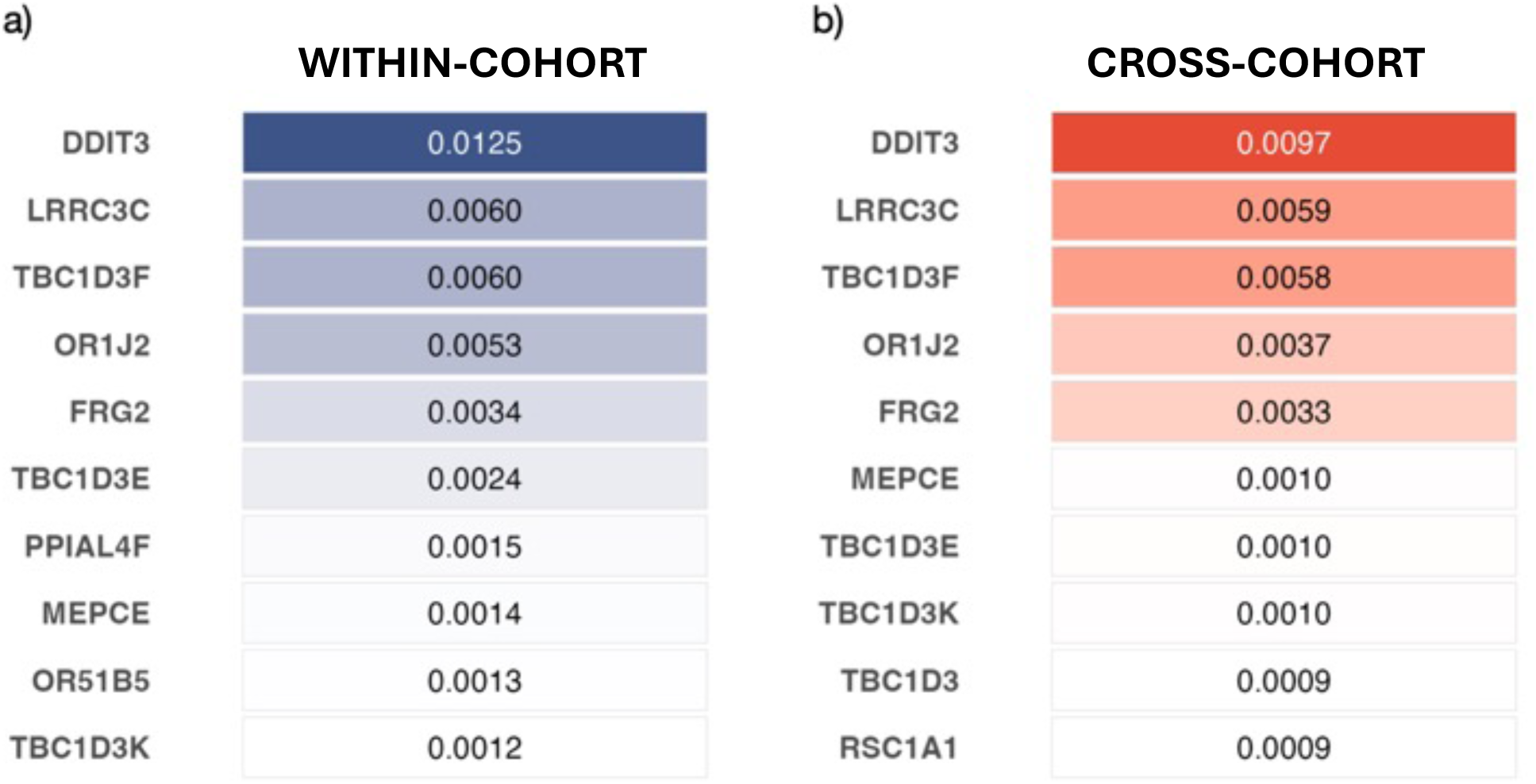
Conserved gene-level attributions from Geneformer embeddings. Ten highest–scoring genes for the frozen Geneformer encoder, computed with Integrated Gradients (PAD baseline; Methods) and aggregated over true-positive test samples, runs/seeds, and outer folds. Panels show the within-cohort (left) and cross-cohort (right) strategies. Values are per-sample positive-only, L1-normalised attribution scores (positive SHAP values kept only), with larger indicates more disease-driving evidence.

**Table 5.** FM approach. Top 20 explainable genes identified via CA-IG for Geneformer embedding model for the within-cohort and cross-cohort approach. SHAP scores normalized using a positive-only approach, then averaged across runs and folds (HGNC symbols). Asterisks (*) represent genes found by both approaches. (Full gene list available in Supplementary Tables S9 (within-cohort) and S10 (cross-cohort)).

| WITHIN COHORT |  |  | CROSS-COHORT |  |  |
| --- | --- | --- | --- | --- | --- |
| GeneID | CA-IG score | Gene name | GeneID | CA-IG score | Gene name |
| **DDIT3\*** | 0.01254973 | DNA Damage Inducible Transcript 3 | **DDIT3\*** | 0.00966168 | DNA Damage Inducible Transcript 3 |
| **LRRC3C\*** | 0.00604412 | leucine rich repeat containing 3C | **LRRC3C\*** | 0.0058623 | leucine rich repeat containing 3C |
| **TBC1D3F\*** | 0.00598382 | TBC1 domain family member 3F | **TBC1D3F\*** | 0.00584974 | TBC1 domain family member 3F |
| **OR1J2\*** | 0.00528166 | olfactory receptor family 1 subfamily J member 2 | **OR1J2\*** | 0.00370744 | olfactory receptor family 1 subfamily J member 2 |
| **FRG2\*** | 0.00337272 | FSHD region gene 2 | **FRG2\*** | 0.00330829 | FSHD region gene 2 |
| **TBC1D3E\*** | 0.00240315 | TBC1 domain family member 3E | **MEPCE\*** | 0.00101443 | methylphosphate capping enzyme |
| **PPIAL4F\*** | 0.0015164 | peptidylprolyl isomerase A like 4F | **TBC1D3E\*** | 0.00099277 | TBC1 domain family member 3E |
| **MEPCE\*** | 0.00144943 | methylphosphate capping enzyme | **TBC1D3K\*** | 0.0009618 | TBC1 domain family member 3K |
| **OR51B5** | 0.00127887 | olfactory receptor family 51 subfamily B member 5 | **TBC1D3\*** | 0.00094298 | TBC1 domain family member 3 |
| **TBC1D3K\*** | 0.00124146 | TBC1 domain family member 3K | **RSC1A1\*** | 0.00092813 | regulator of solute carriers 1 |
| **TBC1D3C\*** | 0.00122356 | TBC1 domain family member 3C | **PPIAL4F\*** | 0.00089612 | peptidylprolyl isomerase A like 4F |
| **RABGAP1\*** | 0.00120914 | RAB GTPase activating protein 1 | **RABGAP1\*** | 0.00089356 | RAB GTPase activating protein 1 |
| **RSC1A1\*** | 0.00119487 | regulator of solute carriers 1 | **DERPC** | 0.00088404 | DERPC proline and glycine rich nuclear protein |
| **TBC1D3\*** | 0.00118897 | TBC1 domain family member 3 | **SPDYE14\*** | 0.00075791 | speedy/RINGO cell cycle regulator family member E14 |
| **TBC1D3H\*** | 0.00116252 | TBC1 domain family member 3H | **TBC1D3C\*** | 0.00074984 | TBC1 domain family member 3C |
| **SPDYE10\*** | 0.00109991 | speedy/RINGO cell cycle regulator family member E10 | **PCDHGC5\*** | 0.00073012 | protocadherin gamma subfamily C, 5 |
| **SPDYE15** | 0.00100317 | speedy/RINGO cell cycle regulator family member E15 | **TBC1D3H\*** | 0.0007123 | TBC1 domain family member 3H |
| **PCDHGC5\*** | 0.00097082 | protocadherin gamma subfamily C, 5 | **PIGY** | 0.00069504 | phosphatidylinositol glycan anchor biosynthesis class Y |
| **PARP2\*** | 0.00096208 | poly(ADP-ribose) polymerase 2 | **PARP2\*** | 0.0006938 | poly(ADP-ribose) polymerase 2 |
| **SPDYE14\*** | 0.00093309 | speedy/RINGO cell cycle regulator family member E14 | **SPDYE10\*** | 0.00068962 | speedy/RINGO cell cycle regulator family member E10 |

The five most predictive genes across both settings were *DDIT3* (DNA Damage-inducible transcript 3-encoding gene), *LRRC3C* (Leucine Rich Repeat Containing 3C-encoding gene), *TBC1D3F* (TBC1 Domain Family Member 3F), *OR1J2 (*Olfactory Receptor Family 1 Subfamily J Member 2-encoding protein) and *FRG2 (*FSHD Region Gene 2-encoding protein) (Fig 4).

The most important gene for endometriosis disease prediction using both the within-cohort and cross-cohort approaches was *DDIT3*, which encodes a transcription factor which mediates response to cellular stress, with a main role in promoting apoptosis during conditions of severe stress. Endoplasmic reticulum (ER) stress is being increasingly recognized as playing a major role in the pathogenesis of endometriosis (27,28), and DDIT3/CHOP is a crucial transcription factor activated by ER stress which has also been shown to further upregulate ER stress and apoptosis in endometriosis disease (29). DDIT3 promotes apoptosis (30), which may drive the degeneration of ectopic tissue. *DDIT3* has been strongly linked to a variety of other diseases including inflammatory diseases, cancer, neurodegenerative diseases and metabolic diseases (31,32). Its’ overexpression has been correlated with poor breast cancer prognosis and implicated in disease progression via its role in cellular stress response pathways (33). *DDIT3* plays key roles in inflammatory diseases by activation of several inflammatory pathways including NF-kb pathway (34) and is upregulated by cytokines TNF-a and IL-1B. *DDIT3* overexpression has also been linked to reduced antiviral innate immune activity, encouraging viral replication and disease (35). This gene is a strong candidate for further exploration and validation.

The second ranked gene, *LRRC3C,* encodes a (predicted) membrane-located protein containing leucine-rich repeats motifs (LRR), important for protein-protein interactions. Although mechanistic studies linking *LRRC3C* with disease remain limited, *LRRC3C* has been implicated as important in inflammatory disease, particularly in inflammatory bowel disease susceptibility via its impact on cell proliferation and apoptosis (36).

The protein encoded by the third most important gene *TBC1D3F* is one of eight paralogs (A-H) belonging the TBC1D3 family. Members of this family of molecules have been found to be over expressed in several cancer types (prostate, breast, pancreatic) (37) and have been associated with poor prognosis of kidney renal cell carcinoma (38). *TBC1D3F* was found to be overexpressed in the epithelial ovarian cancer cell lines and primary tumors compared to normal tissues (39). Alongside *TBC1D3F*, several members of the TBC1D3 family are present in the top 20 list of most important genes for endometriosis disease prediction, including *TBC1D3E*, *TBC1D3K* and *TBC1D3RC.* The TBC1D3 locus is frequently associated with cancer, amplified in 15% of prostate cancers, and the TBC1D3 cluster is extremely structurally variable (40,41).

The amplified presence of genes from the TBC1D3 family (6/20 of the topmost important genes for both within and cross-cohort models) makes this family a strong candidate for experimental validation.

Olfactory receptors (ORs) are expressed in several non-olfactory tissues including the endometrium and are linked to inflammation through specific signaling pathways. Although information on OR1J2 is relatively sparse, other ORs, for example, OR5H2, have been identified as a potential target in endometrial cancer biology related to the IGF1 signaling pathway, and may be involved in cancer protection in some individuals (42).

### LLM-driven knowledge graph inference reveals enriched biological processes linked to endometriosis

Having first investigated the individual functional roles of the top genes and their potential relevance endometriosis disease pathology, we next sought to characterise the shared molecular landscape of these genes to determine whether they shared any links or associations with each other and/or with endometriosis. Because the molecular mechanisms of endometriosis remain poorly characterised, we employed a literature-derived knowledge-graph and LLM-inference approach to do this, which may reveal relationships which traditional bioinformatic gene-centric analysis may miss. For each of the five most important genes for endometriosis prediction (*DDIT3*, *LRRC3C, TBC1D3F, OR1J2* and *FRG2*), we systematically retrieved all PubMed-indexed articles in which it was mentioned and identified semantic relationships between that gene and identified biomedical entities such as disease, gene, pathway, proteins etc. All entities and relationships were represented as entity-entity-relation triples and integrated into a unified bipartite knowledge graph.

The resulting knowledge graph contained 312,598 triples linking 81,219 distinct entities of which 9,835 were proteins, 8,669 were diseases, 5,800 were genes, 4,698 were biological process and 1,367 were drugs (Full list entity types in Supplementary Table S11). The graph contains 33,092 distinct relation types with the most common being co-expression (4,910 triples) and regulation (4,095 triples) (Full list relation types in Supplementary Table S12). First, we investigated the similarity of connected entities and relations between the five important genes by calculating i) normalized shared-entity overlap, and ii) relation-style similarity for each gene pair (see Methods). As expected, most gene pairs exhibited relatively low normalized shared-entity overlap, suggesting genes tended to engage with largely non-overlapping entity sets (Supplementary Table S13). Due to the vastly larger number of triples extracted for gene *DDIT3* than the other important genes, 27%-43% of unique entities from genes *LRRC3C, TBC1D3F, OR1J2* and *FRG2* were shared with *DDIT3* (Supplementary Table S13).

To investigate how our five important genes may converge on common biological mechanisms linked to endometriosis disease pathology, we reduced the multi-gene knowledge graph to a compact JSON structure containing shared entities, shared relations and inter-gene similarity score, and provided as structured context to an LLM to suggest biologically plausible mechanisms supported by the evidence. Although the five genes have distinct primary functions, the LLM results highlighted their roles in several convergent biological themes linked to endometriosis. ER stress is being increasingly recognized as playing a major role in the pathogenesis of endometriosis (27,28), and is enhanced by *DDIT3/CHOP* in endometriosis (29). The LLM results correctly characterize *DDIT3*s potential role in endometriosis disease and links it to the role of *TBC1D3F* in vesicle-mediated cytokine release (43), suggesting a shared influence on inflammatory and stress-response pathways in endometriosis. LLM results also suggest a role of *TBC1D3F* in endometriosis pathogenesis via its role in dysregulated vesicle trafficking, which can alter secretion of proinflammatory cytokines and growth factors that promote ectopic endometrial implantation and contribute towards the pro-inflammatory milieu. This is supported by the literature which indicate that extracellular vesicles play a role in endometriosis disease pathogenesis (44). LLM results suggest a joint contribution of genes *LLRC3C* and *FRG2* to endometriosis pathogenesis through processes related to enhanced adhesion and fibrosis, mediated by extracellular-matrix remodeling (*LLRC3C*) and fibroblast proliferation/fibrosis (*FRG2*). In the literature *LRRC3C* is only partially characterized with predicted roles, however LRR (leucine-rich repeat) proteins have been implicated in cell-adhesion and extra-cellular-matrix (ECM) remodeling (45,46). Enhanced adhesion of endometrial stomal cells to peritoneal surfaces and altered ECM are key steps in endometrial lesion establishment (47,48).

### BMFM attribution rankings for GSEA reveals enriched gene pathways linked to endometriosis pathophysiology

To further investigate associations between the model-derived attribution scores and potential endometriosis disease mechanisms, we performed gene set enrichment analysis (GSEA) from the cross-cohort embedding descending gene list ranked by CA-IG score (see Methods). This analysis identified several positively enriched pathways (FDR < 0.05), indicating a significant association with disease classification. Enriched disease pathways were predominantly related to inflammatory cytokine signaling, cell stress and cell survival, mechanisms consistent with the current understanding of endometriosis disease pathophysiology (Supplementary Figure 1, Supplementary Tables S14-S17). Within cytokine signaling, pathways responsible for positive regulation of type II interferon (IFN-gamma), interleukin-17 (IL-17) and interleukin-8 (IL-8) were all significantly enriched amongst top ranking genes (FDR < 0.05, Supplementary Figure 1). Recent evidence has suggested IL-17 may be an important mediator of endometriosis disease, with elevated levels observed across several endometriosis studies (49,50) and sample types including endometriotic lesions (51) and peritoneal fluid (52) from women with endometriosis. Furthermore, higher levels of IL-17-producing cells have been found in the peripheral blood of women with endometriosis, accompanied by increased IL-17 protein expression in their endometriotic tissue (53), indicating both systemic and local involvement of IL-17 in endometriosis disease. IL-8 is also frequently elevated in the peritoneal fluid of women with endometriosis, and *in vitro* studies have demonstrated IL-17 stimulation of IL-8 secretion from endometriotic stomal cells, linking both pro-inflammatory cytokines to sustenance of the pro-inflammatory milieu of endometriosis (54). Elevated levels of type II interferon (IFN-gamma) have also been detected in endometriotic tissue and peritoneal fluid of women with endometriosis, supporting a role for IFN-gamma mediated inflammatory and immune dysregulation in endometriosis disease pathophysiology (55,56), however some studies have reported conflicting results (57). Alongside cytokine signaling, GSEA indicated several enriched pathways associated with disease were linked to cell stress and cell survival mechanisms, including endoplasmic reticulum (ER) stress, unfolded protein response (UPR) and regulation of apoptosis signaling pathways, highlighting altered cell stress and survival programs in endometriosis (Supplementary Figure 1). ER stress and UPR activation have been shown in endometriotic tissue, and is thought to reflect a response of ectopic cells to the pro-inflammatory milieu (28,58,59). Enrichment of pathways regulating ER stress-induced apoptosis is consistent with a dysregulated balance between stress responses and programmed cell death, a process which has been widely implicated in endometriotic lesion persistence via survival of endometrial cells via evasion of apoptosis (60,61). Together with existing literature, these findings support the interpretation that stress-altered regulation of cell survival pathways represent a prominent feature of endometriosis pathophysiology.

### Conclusions

In this study, we leveraged a multi-cohort bulk RNA-seq dataset to systematically assess the benefit of RNA foundation model (FM) embeddings as feature representations for endometriosis disease prediction using a downstream classifier. We evaluated the benefits of these FM-derived features using a cohort aware strategy and found that FM-derived representations consistently and substantially outperformed gene-level TPM features under a cross-cohort evaluation approach, where models were required to generalize to entirely unseen datasets. This setting reflects a common challenge in clinical transcriptomics and biomarker discovery, and our results demonstrate that pre-trained RNA representations improve robustness to cohort heterogeneity.

A central contribution of this work is the introduction of classifier-aligned integrated gradients (CA-IG), a computationally efficient approach for interpreting gene-derived FM embeddings. While IG benefits from a GPU for speed, it substantially lowers the computational burden for interpretability of FMs and makes analyses feasible with modest resources. Applying CA-IG to Geneformer embeddings revealed highly stable gene-level attributions across both cohort-aware strategies, in contrast to TPM-based models where predictive genes varied substantially between cross and within cohort approaches. Embedding models consistently prioritised a conserved set of genes, supporting the notion that pretrained encoders can capture generalisable biological knowledge rather than cohort-specific noise. Pathway-level analyses further revealed convergence on inflammatory cytokine signaling, cell stress and cell survival mechanisms, processes strongly implicated in endometriosis disease pathophysiology.

Some limitations remain within this work. First is the small size of several of the individual cohorts, which increased variance in train/test splits. We addressed this using dataset-aware splits, a grouped-cohort outer CV, a nested inner CV for model selection, and multiple seeds with bootstrap confidence intervals. Second was the imbalanced nature of the full cohort, with comparatively few controls. This imbalance likely contributed to the reduced predictive performance of class 0 compared to class 1 and is a common issue in disease/control classification. Our third limitation was associated with inconsistent reporting of key covariates due to small and varied sample sizes (e.g., menstrual-cycle phase, lesion subtype), which prevented adjustment for potential confounders.

RNA FMs offer an accessible and powerful solution for improving the robustness of disease classification in transcriptomics. Here we demonstrated this for endometriosis, a condition hampered by diagnostic delays and biological heterogeneity, where this CPU-friendly approach provided more generalizable predictions and stable, interpretable biomarkers, laying a practical foundation for future application to other diseases and use-cases, leveraging large-scale biological priors in clinically focused research.

## Methods

### Datasets and pre-processing

We assembled a multi-cohort bulk RNA-seq corpus from the Gene Expression Omnibus (GEO) (20) by searching for human studies of endometrial tissue related to endometriosis and restricting results to expression profiling by high-throughput sequencing. Single-cell, spatial transcriptomic studies, and studies with fewer than 10 samples (pre-filtering) were excluded. We used the GEO identifiers for our selected studies to download the associated transcriptomic data for these studies and firstly, to merge and normalise the gene expression count data (TPM) into a unified data matrix, and then to extract harmonised metadata for the same samples i.e., in this case we used an LLM to yield a label for each sample of endometriosis disease yes (class 1) or control/no (class 0). Labels were derived from the harmonised metadata and then further checked and curated by an expert molecular biologist to remove ambiguous cases (e.g., endometrial cancer, unclear disease status). After filtering, the final corpus comprised 12 studies and 334 samples (259 endometriosis, 75 healthy controls). GEO Series IDs were retained as a grouping variable for subsequent cohort-aware cross-validation. Raw counts were converted to transcripts per million (TPM) using GRCh38 (p13) gene lengths and log-transformed (log(1+TPM)). Gene identifiers were harmonised across cohorts, and all matrices were aligned to a common 17,518-gene universe.

### Fold construction

#### Within-cohort

For the within-cohort analysis, we used a dataset-aware, stratified 3-fold split approach. To ensure each dataset was represented in the training set for each fold, we first selected one sample per GEO series or dataset as the ‘anchor’ sample and pinned all anchors to the training set in every fold. We then applied stratified-*K-*fold (*K*=3, shuffled, fixed seed) to the non-anchor samples only, preserving the global class ratio in each test fold. In fold *k* the training set was the union of all anchors and the non-anchor samples not assigned to the *k*-th test split; the test set was the corresponding stratified *k*-th split. This procedure yields zero sample overlap between train and test, keeps test-set class balance approximately intact, and enforces an in-domain condition in which any series or dataset appearing in test also has at least one sample (its anchor) in train. Fold assignments and per-fold counts are reported in Table 6.

**Table 6.** Fold construction for the within-cohort and cross-cohort folds.

| **Within-cohort Folds** | **Train (n)** | **Test (n)** | **Positive proportion (train)** | **Positive proportion (test)** | **Dataset IDs** |
| --- | --- | --- | --- | --- | --- |
| 1 | 226 | 108 | 0.78 | 0.77 | 1, 2, 3, 4, 5, 6, 7, 8, 9, 10, 11, 12 |
| 2 | 227 | 107 | 0.78 | 0.77 | 1, 2, 3, 4, 5, 6, 7, 8, 9, 10, 12 |
| 3 | 227 | 107 | 0.78 | 0.78 | 1, 2, 3, 4, 5, 6, 7, 8, 10, 11, 12 |
| **Cross-cohort Folds** | **Train (n)** | **Test (n)** | **Positive proportion (train)** | **Positive proportion (test)** | **Dataset IDs** |
| 1 | 237 | 97 | 0.78 | 0.77 | 6,7,9 |
| 2 | 228 | 106 | 0.77 | 0.78 | 5,8,12 |
| 3 | 203 | 131 | 0.78 | 0.77 | 2,4,10,11 |

#### Cross-cohort

For the cross-cohort approach, we constructed balanced grouped-cohort splits with *K*=3 outer folds to match the within-cohort approach. Entire datasets (GEO Series IDs) were assigned to test folds by a constrained random search (1,000 trials; fixed seed) that (i) enforced a minimum test size (at least 60 samples per fold) and (ii) minimised the across-fold standard deviation of the positive fraction and of fold sizes. This resulted in test/train dataset assignments as reported in Table 6.

### Extraction of Foundation Model Embeddings

We evaluated five pretrained encoders spanning bulk and single-cell transcriptomic pretraining: BulkRNABERT (18), scFoundation (17), scGPT (16), Geneformer (15) and BMFM-MLM-RDA (19). These models learn transferable structure from large expression corpora and produce compact, context-aware representations (embeddings) for downstream prediction.

For each sample we performed feature harmonisation as follows: (i) mapping the TPM vector to the encoder’s gene vocabulary (e.g. HGNC to Ensembl); (ii) dropping genes absent from that vocabulary; (iii) consolidating duplicate symbols by averaging their TPM values; and (iv) constructing model-specific feature sets (tokens). To extract embeddings per sample, we then executed a single forward pass with the encoder in evaluation mode (frozen weights) and obtained a fixed-dimensional embedding per sample. Embeddings were computed once per sample and cached before any model selection; identical splits and seeds were reused across models for paired comparisons. We intentionally avoided additional downstream scaling or feature selection on embeddings to preserve the pretrained geometry. When multiple HGNC symbols mapped to the same Ensembl identifier, we averaged their TPM values by grouping them by their shared Ensembl identifier prior to input construction. We chose this approach as averaging preserves scale and avoids artificial inflation for genes with many aliases, which could otherwise bias downstream models. Model-specific approaches are detailed below:

#### Geneformer

Geneformer is a transformer pretrained on 104M single-cell transcriptomes with rank–value encodings and masked-token objectives (15). We mapped TPM(log+1) matrices to Ensembl IDs, tokenised with the official toolkit, and extracted the penultimate layer. Mean pooling excluded special tokens (CLS/EOS, PAD) producing a 768-dimension embedding per sample.

#### BulkRNABERT

This encoder-only model was pretrained on bulk RNA-seq (GTEx/ENCODE/TCGA) with a masked-LM objective over discrete expression tokens and permutation-invariant gene order (18). TPM(log+1) matrices were harmonised to the accepted Ensembl set and ordered to the model list. We extracted embeddings from the final layer mean-pooled across tokens to yield a 256-dimensional embedding per sample.

#### scFoundation

ScFoundation is an asymmetric encoder–decoder pretrained on >50 million human single-cell profiles (62). TPM(log+1) vectors were aligned to the 19,264-gene vocabulary (missing genes padded with zeros). The official routine returns a 3,072-dimensional vector comprising four 768-dimensional pools (1: last token, 2: penultimate token, 3: max pooled, 4: mean pooled). We split this file and used the last 768 dimensions for all further analysis (mean-pooled 768-dim block).

#### scGPT

ScGPT is a generative transformer pretrained on 33 million single-cell profiles with masked-attention objectives, jointly learning gene and cell embeddings (16). TPM(log+1) matrices were aligned to the model vocabulary and embedded, yielding a 512-dim vector per sample.

#### BMFM-MLM-RDA

We used the MLM-RDA variant of BMFM: a BERT-base encoder, pretrained with read-depth–aware (RDA) down sampling pretrained via expression masking and RDA downsampling (https://huggingface.co/ibm-research/biomed.rna.bert.110m.mlm.rda.v1, (19)). We used TPM (log+1) data for a single forward pass through the frozen embedder; the final-layer [CLS] vector (768-dimensional) served as the sample embedding.

### Explainable Machine Learning workflow

#### Train and test protocol

We used the open-source explainable-AI tool AutoXAI4Omics (https://github.com/IBM/AutoXAI4Omics) to run a robust classical explainable ML analysis on both TPM and embedding inputs for both the within-cohort and cross-cohort strategies, ensuring consistency, transparency, and reproducibility across model comparisons (63). We chose the AdaBoostClassifier model as our downstream light-weight classifier (64).

We evaluated two input data types within the same AutoXAI4Omics classical-ML pipeline for endometriosis classification: (i) log TPM values and (ii) foundation-model (FM) embeddings derived from log TPM values. We adopted two strategies; i) within-cohort, where train and test contain samples from the same GEO dataset ID, and ii) cross-cohort, where the held-out test fold comprised unseen datasets not represented within the training set. The outer loop used K=3 folds; in each outer fold, models were trained on K-1 folds and evaluated once on the held-out fold. Within the fold’s training set, hyperparameters were tuned by stratified 5-fold cross-validation, selecting the configuration that maximised weighted-F1 (primary metric), then refitting that configuration on the full training data before testing. To assess robustness, each outer split was repeated with five different random seeds; we report means across folds and seeds. All preprocessing (scaling, feature transforms, encoders and feature selection) was fitted strictly on the training side and applied unchanged to validation/test to avoid leakage.

#### TPM data preparation for ML

For all AutoXAI4Omics runs with TPM data we performed feature selection using the following two inbuilt methods; (i) first a variance filter removed near-constant genes (threshold = 1); and (ii) secondly a univariate ANOVA test (F-statistics) was used to select the top *k* genes, where *k* was chosen automatically from a base-10 grid with a minimum of 25 features (no explicit maximum). For each candidate *k,* a small random forest was trained on the selected genes and evaluated by F1 score. From the set of values yielding high performance, we selected the smallest *k,* prioritising a smaller feature set while retaining performance.

#### FM-embedding data preparation

For all FM vectors we disabled standardisation and feature selection, treating embeddings as fixed representations to avoid distorting pretrained geometry.

### Post-AutoXAI4Omics Bootstrapping and significance calculation

To obtain a robust and sound estimate of all models performance variability and allow statistically valid comparison of models, we calculated confidence intervals for fold-averaged metrics via hierarchical bootstrapping, with 10,000 iterations. For each model we had performance metrics from 3 folds and 5 seed runs per fold, totaling 15 runs per model. Each bootstrap iteration consisted of drawing 3 folds with replacement, and per selected folds drawing 5 seed runs with replacement. For each metric ‘fold means’ were calculated which were then used to calculate ‘overall means’ across all folds. The same approach was used to calculate the effect size (while preserving paired seeds across models), defined as the difference in model metrics to the baseline model (TPM input, not embeddings). Bootstrap estimates of the mean and effect size over 10,000 iterations were used to construct 95% confidence intervals. To test whether performance of paired models differed significantly we leveraged the same nested-bootstrap samples that we used for confidence intervals. In each of the 10,000 bootstrap iterations we paired the corresponding runs of two models (same index), computed the difference in performance metric for each fold, and averaged these differences over the 3 folds. This yielded one bootstrap difference value per metric per iteration. Under the null hypothesis that the models have equal performance, the distribution of differences would be centred on zero (no difference). We therefore we calculate two-tailed p-values as follows: i) compute the proportion of bootstrap differences which lie above zero, and the proportion that lie below zero, ii) take the smaller proportion (representing the least likely direction of difference) and iii) we double this to get a 2-tailed p-value. This approach was applied to all pairwise comparisons (baseline/vanilla vs. all BMFMs) for the metrics F1-score, F1-score (class 0), F1-score (class 1) and accuracy.

### Evaluation Metrics

Primary reported metrics were weighted-F1, F1_class0_, F1_class1_ and accuracy with 95% CIs obtained via bootstrapping as described above. To characterize the benefit of frozen FM embeddings over the baseline TPM we report paired effect sizes for weighted-F1, F1_class0_, F1_class1_ and accuracy, computed paired by seed within fold and summarized with the same bootstrap (paired 95% CIs).

### Explainability

#### SHAP-based gene attribution (TPM models)

We used Shapley Additive exPlanations (SHAP) [ 26, 27] as implemented in AutoXAI4Omics to quantify per-gene contributions to the positive-class F1 score. We used the TreeSHAP explainer for the AdaBoost classifier, fitted on training folds only and applied to the held-out test split. To focus on disease-driving signal (rather than model error or protective evidence), we collected and aggregated SHAP scores from all folds (n=3) and all runs (n=5) of the AdaBoost classifier only for **true-positive** test samples (ground-truth is ‘disease’ and predicted positive threshold >= 0.5). To allow comparison of SHAP scores across samples, aggregated true-positive SHAP scores were subject to a positive-only normalization, where positive SHAP scores were normalized per sample by scaling each sample’s positive SHAP vector into a distribution summed to 1(only positive SHAP scores were retained to allow focus on features driving disease prediction) Normalized SHAP values were then averaged across folds and seeds to generate a normalized average SHAP value per feature.

#### Geneformer attribution (classifier-aligned integrated gradients)

Focusing on our best performing FM Geneformer, in this study we used transcriptomic information as input into the FM to derive embeddings (see Methods: Extraction of FM embeddings, Geneformer). We then used these embeddings to train a downstream classifier (AdaBoost) to predict endometriosis status, but notably this classifier was not fine-tuned jointly with Geneformer. Using explainability techniques, our goal was to trace disease predictions back through the FM encoder to identify which input genes were most responsible for the classification decision from our resultant AdaBoost model. To derive this explainability, we developed an approach that we call classifier-aligned integrated gradients (CA-IG) to attribute the FM token inputs (gene tokens) to the downstream disease score used for classification using Integrated Gradients (IG), a well-established attribution method (65). However, the “classifier-aligned” aspect of our approach, computing gradients using SHAP values rather than through the entire model, brings novelty and utility, dramatically reducing computational cost while maintaining gene-level information about what drives predictions.

When a tokenised RNA-seq sample is processed by Geneformer, it produces a matrix of dimension *L* × *D*_*m* (where L represents length and D_m is the embedding size). The Geneformer encoder at layer ℓ (the penultimate layer in all our experiments) transforms this into hidden states of dimension L×D. We focused on real tokens only, excluding special tokens like CLS, EOS, and PAD, and applied a masking procedure to obtain the sample embedding. Therefore, the sample embedding, denoted as *z*(*X*), was created by mean-pooling the masked hidden states over all real tokens, weighted by their mask values *m* ∈ {0,1}^*L*. This produced a D-dimensional vector (768 dimensions in this instance) that represented the entire sample in the embedding space.

Our downstream classifier AdaBoost operates on this D-dimensional embedding space to produce a positive-class score (the disease probability). Since the classifier is not fine-tuned jointly with Geneformer we used a sample-specific surrogate direction derived from the realised SHAP attribution vector of the classifier’s positive logit with respect to this embedding. This surrogate direction 𝑤(*x*) exists in the same D-dimensional space as the embedding. We then defined a scalar objective 𝑠(*X*) as the inner product between this SHAP direction and the sample embedding *z*(*X*). This scalar objective is locally faithful to the classifier and enabled efficient backpropagation into the token embeddings without requiring differentiation through the separate classifier. We call this approach classifier-aligned integrated gradients (CA-IG) because it aligns the gradient direction with the classifier’s behaviour.

Computationally, this approach converts the vector sensitivity into a single directional derivative along the SHAP direction 𝑤(*x*). Using the linearity of gradients, computing the gradient of this scalar objective with respect to the embedding X can be expressed as a weighted sum of per-dimension gradients, where each dimension is weighted by the corresponding SHAP weight. This means a single backward pass per IG step yields the same weighted gradient that a user would obtain by computing per-dimension gradients separately and weighting them afterward. This reduces the computational cost from 𝑂(𝑆 · *D*) to 𝑂(𝑆) backpropagations for S integration steps across embedding dimension D. To prevent data leakage, the SHAP direction 𝑤(*x*) is computed using only the training fold, then applied to held-out test samples.

IG is a path-based attribution method that accumulates gradients along a straight-line path from a baseline to the actual input. We used a PAD baseline B of dimension *L* × *D*_*m*, where each position was set to the learned pad token vector. The IG from baseline B to input X were calculated by integrating the partial derivative of the objective with respect to the input along the path from B to X, parameterised by α from 0 to 1. This continuous integral was approximated using a Riemann sum with S=32 discrete steps along the path. At each step, we evaluated the gradient at a point that interpolated between the baseline and the actual input. Token-level scores were obtained by summing these IG across all channels (the D_m embedding dimensions) for each token position, resulting in a score 𝑎_𝑖^ + (*x*) for each gene token i. These token scores were then mapped to genes by removing special tokens and using the tokeniser’s gene dictionary to form gene-level attributions *g*^ ± (*x*).

Akin to the SHAP analysis, we focused on disease-driving signals by reporting summaries that emphasised positive (disease-promoting) contributions. For each sample x, we normalised the attributions per sample before aggregating. We created a positive-only summary by taking the maximum of each gene’s attribution and zero, then normalising these positive values to sum to 1 (with a small constant ε added to the denominator for numerical stability). This gave us ĝ_𝑖^ + (*x*), representing gene i’s relative positive contribution to the disease prediction for sample x. We also computed a signed summary ĝ_𝑖(*x*) that preserved the direction of effects by normalising using the sum of absolute values across all genes, again with the stabilising constant ε. We attributed only true-positive patients—specifically, test samples where the ground truth y=1 (disease present) and the predicted probability exceeded 0.5. For outer CV fold k and repeat r, we identified the set 𝑇_𝑘, 𝑟 of true-positive test samples. We then averaged the normalised attributions within each run, within each fold, and finally across all folds.

The final importance score 𝐼𝐺𝐼*m*𝑝_𝑖^ + for gene i was calculated by averaging over K folds, R repeats, and all true-positive samples within each fold-repeat combination, using the positive-only normalised scores. Similarly, 𝐼𝐺𝐼*m*𝑝_𝑖^ ± used the signed normalised scores to capture directional effects. This multi-level averaging produced stable, generalisable gene importance scores that reflect consistent patterns across different data splits and random initialisations, minimising the influence of any single model or sample. To our knowledge, CA-IG is the first attribution protocol tailored to frozen RNA foundation-model embeddings with a separate downstream classifier, and it reduces the per-sample IG complexity from O(S·D) to O(S) while remaining CPU-efficient. We report IGImp+ in the main text and provide IGImp± in the Supplementary material.

### GSEA from attribution ranks

To identify enriched biological pathways associated with disease we performed gene set enrichment analysis (GSEA) using pre-ranked gene lists derived from feature importance scores. Four separate analyses were performed with results from the following approaches: a) within-cohort TPM model and associated gene SHAP scores; b) cross-cohort TPM model and associated gene SHAP scores; c) within-cohort FM embedding model (Geneformer) and associated gene attribution scores from CA-IG and d) cross-cohort FM embedding model (Geneformer) and associated gene attribution scores from CA-IG. For TPM-based analyses gene importance scores were obtained from SHAP values of the disease classification model whilst FM GSEA analyses were run with gene attribution scores calculated from the classifier-aligned integrated gradients method. The gene universe was defined as all genes present in the input TPM matrix. For each of the four analyses, gene importance scores were mapped to the gene universe and genes were ranks in descending order of their signed importance. Pre-ranked GSEA was performed using GSEApy (66) with 10,000 permutations, testing using gene sets from the Enrichr GO Biological Processes library (67). Pathways significantly enriched at either end of the ranked gene list were identified at FDR < 0.05, with positive normalized enrichment score (NES) indicating enrichment among disease-associated genes and negative normalized enrichment score (NES) indicating enrichment among healthy-associated genes. Parameters used are as follows: min_size= 15, max_size= 500, permutation_num= 10,000, Benjamini–Hochberg FDR control (seed= 42). For each term we report the NES, nominal P value, and FDR-corrected P-value. “Leading-edge fraction” denotes the leading-edge size divided by the gene-set size.

### Knowledge graph creation and LLM inference

To identify unexplored relationships between our most explainable genes and endometriosis in a sparsely characterised research area, we applied a literature-driven knowledge graph and large language model-based inference method to systematically integrate and reason over published evidence. This yielded a literature-grounded representation of gene-endometriosis relationships suitable for potential hypothesis generation. Each candidate gene was treated as an independent query entity and used to retrieve all PubMed-indexed articles in which it was mentioned, using open-source software DeepSearch (https://ds4sd.github.io/). From the resulting corpus, we extracted paragraph-level text segments containing co-occurrences of each entity and the endometriosis. These texts were processed using named entity recognition and relation extraction to identify biomedical entities such as genes, diseases, proteins, pathways, and used to identify the semantic relationships amongst them. All entities and relationships were represented as entity-entity-relation triples, and integrated into a unified knowledge graph, with literature provenance retained at document and paragraph levels. All inferred associations were constrained by the underlying graph structure and traceable to one or more publications. To enable LLM-based graph-enabled inference to identify higher-order or implicit relationships supported by the literature and interpret the shared relational patterns in the compact KG, we supplied a compact JSON summary to a locally deployed LLM (GPT-20b) executed using Ollama inference engine. We also calculated relation-style similarity and shared-entity overlap for all our gene pairs (see below) and provided that to the LLM to aid inference.

#### Relation-style similarity (weighted Jaccard similarity)

To quantify similarity of connected entities and relations between the five important genes in the knowledge graph, we calculated a weighted Jaccard similarity for each pair of important genes, quantifying the similarity of their relation-type profiles across all shared entities. For each pair of important genes and for each entity common to both genes, we calculated an intersection weight by summing the minimum frequency observed for each relation type, as follows:

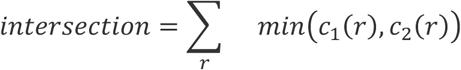

Next, for that specific entity we captured the union weight which measures total combined relation usage, so taking the maximum of the two counts as follows:

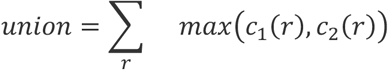

We then calculate the Jaccard similarity (intersection/union) with 1 indicating that the two genes share exactly the same relation types (for shared entities) with same frequencies, and 0 indicating the two genes share no common relation types for any shared entities.

#### Normalized shared entity overlap

For each gene pair, to account for substantial variation in the number of triples per gene due to varied representation in the literature, we also calculated shared entity overlap directionally as the proportion of shared entities from gene1 that were also present in gene2, and reciprocally from gene2 to gene 1. This normalization allows overlap to be evaluated relative to each genes individual representation in the corpus.

#### Shared-entity overlap (Jaccard similarity)

To complement the relation-style similarity we also calculate a shared-entity Jaccard to quantify shared entities between each gene pair. For each gene pair, this is defined as the proportion of shared (overlapping) entities relative to the total number of distinct entities linked to each gene. For two genes *g*_1_ and *g*_2_, the shared-entity jaccard is calculated as follows:

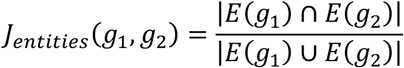

## Declarations

### Ethics approval and consent to participate

Not Applicable – all data used in this study was publicly available from the Gene Expression Omnibus (GEO)

### Consent for publication

Not Applicable

### Availability of data and materials

All datasets used in this study are bulk RNA-seq data that is publicly available, **Table 1** gives a summary of the cohorts used in this study, detailing Gene Expression Omnibus (GEO) ID’s. The foundation models and AutoXAI4Omics used in this study are all open source (see citations in text). However, we describe a new interpretability method for the FM embeddings that we call CA-IG. We will release this code as an open source repo (at https://github.com/IBM/) upon acceptance of publication.

### Authors’ Contributions

LJG & JK designed the study, LJG/JK/RT/MA/SC/AE contributed towards workflow conceptualization, LJG, JK & NM performed the AI technical analysis, NM, JK & LJG performed the KG technical work with contributions from JB, NC created the CA-IG methodology supervised by LJG & JK, JK, LJG & NM wrote and edited the manuscript, MM contributed to BMFM embedding extraction, transcriptomic data collection and curation, all authors reviewed code and/or manuscript.

## Acknowledgements

This work was supported by the Hartree National Centre for Digital Innovation, a collaboration between STFC and IBM.

