## Supplementary Figure 1 for "RNA foundation models enable generalizable endometriosis disease classification and stable gene-level interpretation"

### Supplementary Material

#### Supplementary Figures 1

#### Supplementary Tables 1-17 – see additional file

### Supplementary Figures

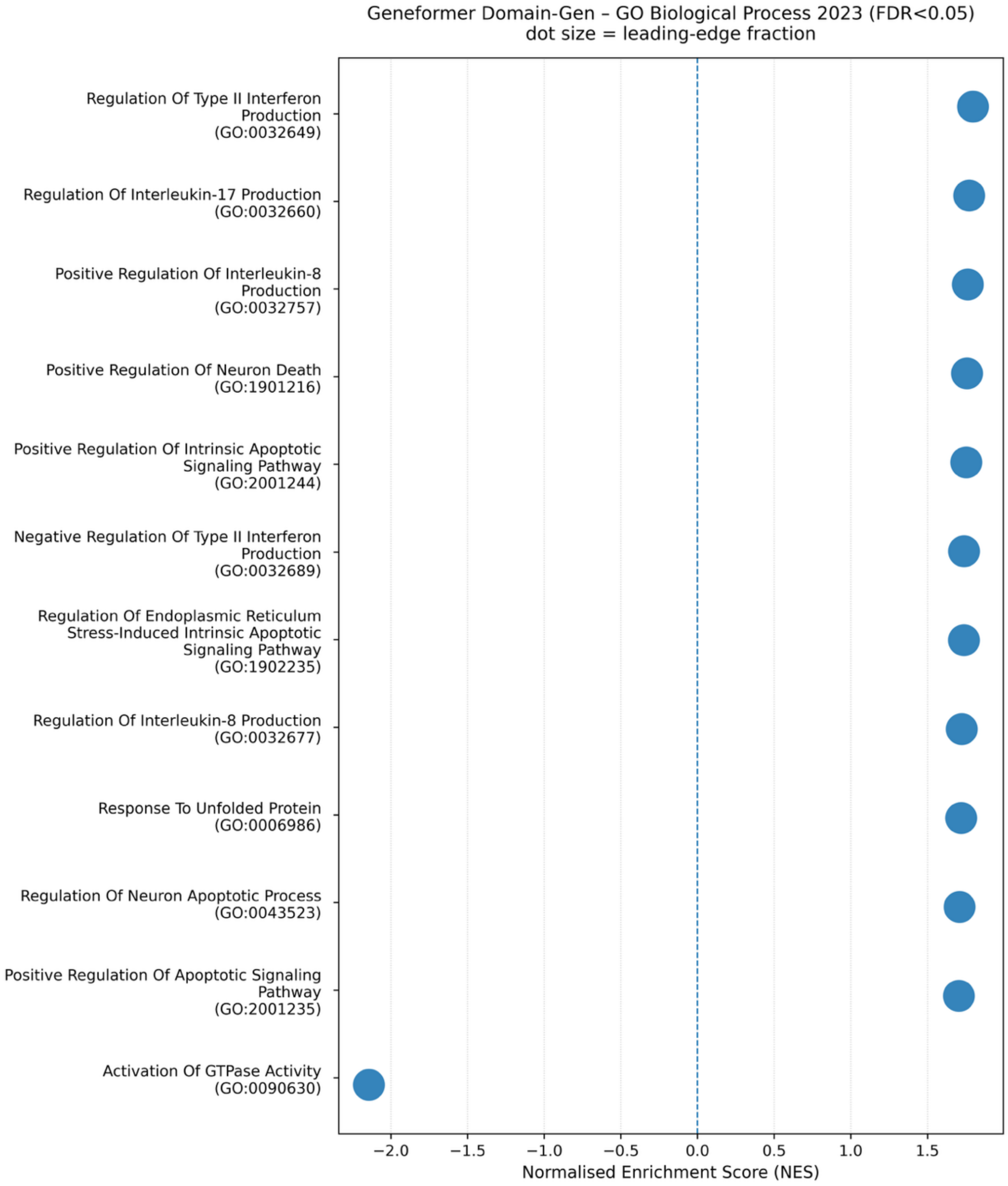


Supplementary Figure 1. Gene set enrichment analysis (GSEA) dot plot of disease-associated pathways identified from cross-cohort FM embedding gene attribution scores. Genes were ranked in descending order of signed importance scores derived from CA-IG of cross-cohort results. Pre-ranked GSEA was performed using the Enrichr GO Biological Processes gene set library with 10,000 permutations. Only pathways significantly enrihced at an FDR < 0.05 are shown.
